# Validity of the cell-extracted proteome as a substrate pool for exploring phosphorylation motifs of kinases

**DOI:** 10.1101/2023.03.20.533483

**Authors:** Tomoya Niinae, Naoyuki Sugiyama, Yasushi Ishihama

**Affiliations:** Graduate School of Pharmaceutical Sciences, Kyoto University, Kyoto 606–8501, Japan; Laboratory of Clinical and Analytical Chemistry, National Institute of Biomedical Innovation, Health and Nutrition, Ibaraki, Osaka 567-0085, Japan

## Abstract

Three representative protein kinases with different substrate preferences, ERK1 (Pro-directed), CK2 (acidophilic), and PKA (basophilic), were used to investigate phosphorylation sequence motifs in substrate pools consisting of the proteomes from three different cell lines, MCF7 (human mammary carcinoma), HeLa (human cervical carcinoma), and Jurkat (human acute T-cell leukemia). Specifically, recombinant kinases were added to the cell-extracted proteomes to phosphorylate the substrates *in vitro*. After trypsin digestion, the phosphopeptides were enriched and subjected to nanoLC/MS/MS analysis to identify their phosphorylation sites on a large scale. By analyzing the obtained phosphorylation sites and their surrounding sequences, phosphorylation motifs were extracted for each kinase-substrate proteome pair. We found that each kinase exhibited the same set of phosphorylation motifs, independently of the substrate pool proteome. Furthermore, the identified motifs were also consistent with those found using a completely randomized peptide library. These results indicate that cell-extracted proteomes can provide kinase phosphorylation motifs with sufficient accuracy, even though their sequences are not completely random, supporting the robustness of phosphorylation motif identification based on phosphoproteome analysis of cell extracts as a substrate pool for a kinase of interest.

## Introduction

Protein phosphorylation plays an important role in intracellular signal transduction and regulates various biological functions such as cell growth and differentiation. Phosphoproteomics using mass spectrometry (MS) has made it possible to identify thousands of phosphorylation sites in a single measurement (Bekker-Jensen et al., 2020; Humphrey et al., 2015). However, despite the vast amount of phosphorylation site information stored in public databases (Dinkel et al., 2011; Hornbeck et al., 2015; Keshava Prasad et al., 2009), little is known about the protein kinases associated with individual phosphorylation sites (Needham et al., 2019).

Attempts have been made to obtain information on kinase-substrate relationships *in vivo* by perturbing kinase activity in living cells through kinase inhibitor treatment or knockdown/knockout of specific kinases (Imami et al., 2012; Pan et al., 2009). However, these methods also indirectly affect downstream kinases, thus hindering direct substrate identification (Bodenmiller et al., 2010). Alternatively, *in vitro* kinase assays have been used to validate direct interactions between kinases and candidate substrates. Although the information obtained by this *in vitro* approach does not necessarily reflect *in vivo* phosphorylation events, *in vitro* kinase assay is the simplest and easiest way to obtain reliable site information, because phosphorylation motifs are not expected to change significantly between *in vitro* and *in vivo* conditions. Therefore motif information obtained from *in vitro* kinase reactions is one of the most important filters to narrow down *in vivo* substrate candidates.

Traditionally, motifs from *in vitro* kinase assay have been obtained from random sequence libraries of peptides immobilized on supports (Hutti et al., 2004; Miller et al., 2008). Realistically, however, there is a limit to the number of combinations of amino acid sequences that can be examined (for example, 10 positions around the phosphosite in total, with 20 amino acid options per position, would result in a total of 20^10^ combinations), So, practical evaluation has to be done using mixtures (Hutti et al., 2004; Johnson et al., 2023; Miller et al., 2008). Another issue is that the influence of immobilization of substrate peptides cannot be eliminated (Inamori et al., 2005, 2008). On the other hand, by using samples extracted from cells as a substrate pool (Huang et al., 2007), thousands of substrates can be easily identified by MS (Imamura et al., 2014; Sugiyama et al., 2019). However, it has not been established how the proteome profile of the cell-extracted substrate pool used affects the motifs extracted, or whether there is a difference between the motifs identified from cell-extracted substrate pools with human proteome bias and from random sequence peptide libraries without such bias.

In this study, three kinases with acidophilic, basophilic, and proline-directed phosphorylation motifs were used as models, and phosphorylation motifs from three different cell line-derived substrate pools were compared. Motifs from a random sequence peptide library were also compared to determine whether phosphorylation motifs could be extracted from the cell-line-derived substrate pools without bias.

## Results

### *In vitro* kinase substrates obtained from different cell lines

The position weight matrix (PWM) (Wasserman & Sandelin, 2004) represents the characteristics of the sequence around the phosphorylation site targeted by a given kinase (Imamura et al., 2017). The columns of the matrix represent the positions relative to the phosphorylation site, and the rows represent the amino acids. The values of the matrix are log2 transformed probabilities that a given amino acid occurs at a given position. These probabilities have been used to predict putative substrates, and the PWM scores represent the certainty of those sequences as substrates (see Methods). This approach has been implemented in several kinase substrate prediction methods and also applied to *in vivo* kinase substrate identification (Imamura et al., 2017).

The construction of PWMs requires substrate information on phosphorylation sites and their surrounding sequences, which can be obtained very efficiently by analysis of *in vitro* kinase reactions on proteins extracted from cells. However, when proteins are extracted from different cell types, the proteomic profiles are different, and thus the resulting in vitro substrate peptides may differ depending on the cells used. The impact of this difference on the PWM is not known.

Here, we assessed whether PWMs based on *in vitro* substrates obtained from cell-extracted proteins are independent of the type of cells used to acquire the *in vitro* substrates. We selected three cell lines (HeLa, MCF7, and Jurkat) that are commonly used and that are known to have different proteome profiles (Geiger et al., 2012). For the *in vitro* kinase reaction, we selected three kinases with different substrate specificities (CK2: acidophilic, PKA: basophilic, and ERK1: proline-directed). Proteins were extracted from each cell line and TSAP was added to dephosphorylate the endogenous phosphorylation sites. TSAP was inactivated by heating, and then the recombinant kinase was added to the substrate pool proteins. The proteins were digested with trypsin, and the phosphorylated peptides were enriched by HAMMOC and analyzed by LC/MS/MS (Sugiyama et al., 2007). The phosphorylation sites quantified in the control sample without spiked kinase (kinase (-)) were excluded from those identified in the sample that reacted with each kinase (kinase (+)), and the final list of phosphorylation sites generated by the *in vitro* kinase reaction was determined (Table S1a, S1b, S1c, S2). To confirm that the sets of in vitro substrate peptides derived from the different cell-extracted substrate pools were different, we compared the MS signal intensities (Fig. 1a, S1a). The correlation of signal intensities of substrate peptides in triplicate experiments was high for each kinase (0.90 < R < 0.98), while the correlation of signal intensities for substrate peptides derived from different cell lines was low (0.64 < R < 0.79). Furthermore, when the signal intensity ratios of proteins and those of in vitro substrates between pairs of cell lines were plotted, the intensity ratios of proteins and in vitro substrates were correlated (Fig. 1b, S1b, Table S1d). These results indicate that the in vitro substrate profile is influenced by the proteome expression profile in each cell line.

**Fig. 1.**
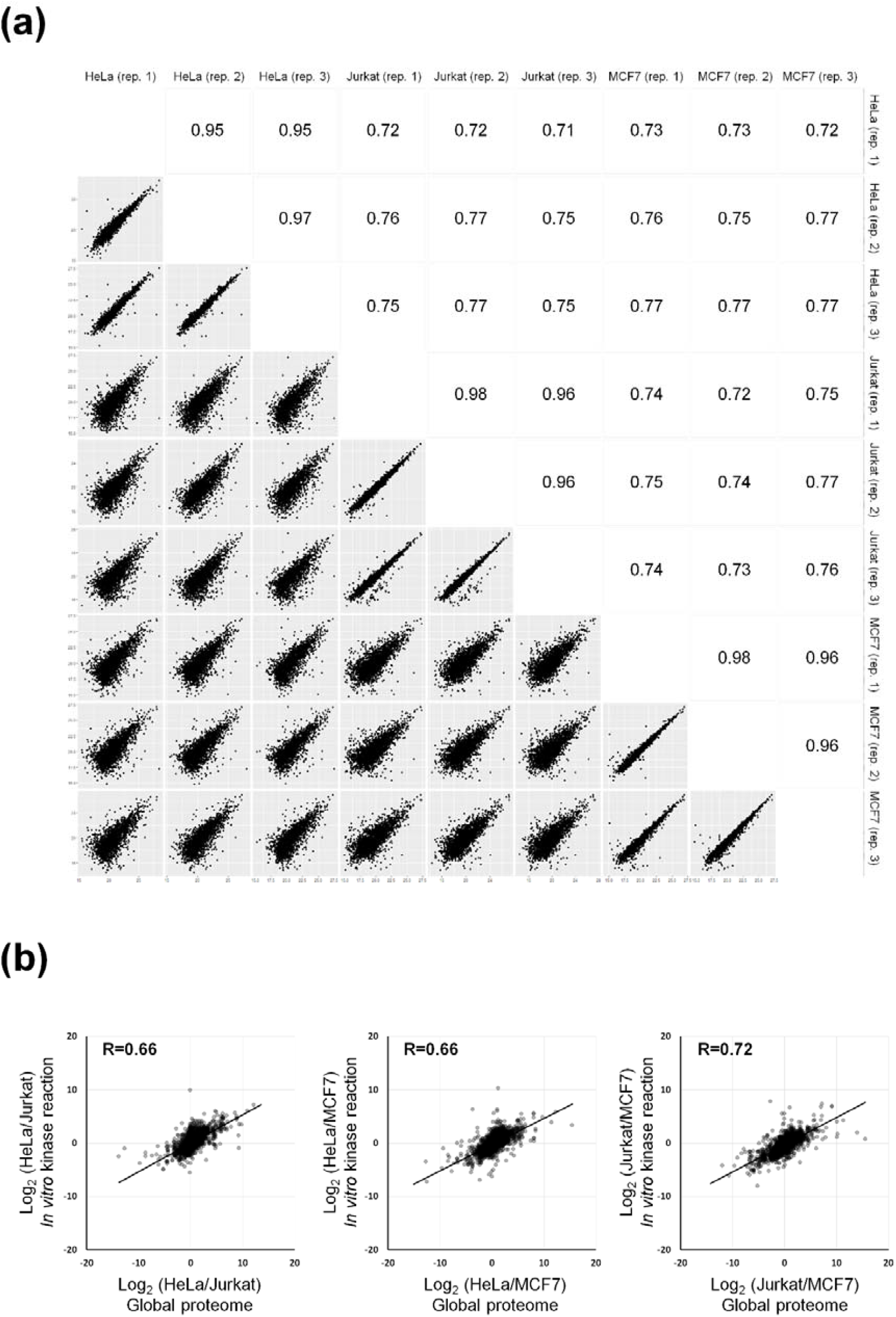
Relationship between *in vitro* ERK1 substrates obtained and protein expression levels. **(a) Correlations of signal intensities of *in vitro* ERK1 substrates derived from triplicate experiments with each of the three cell lines** Pearson correlation coefficients of signal intensity (log 2) were calculated for phosphorylation sites (pS, pT) obtained from triplicate *in vitro* kinase reactions for each set of cell-extracted proteins. **(b) Correlations between protein expression levels and *in vitro* ERK1 substrate abundance** The normalized ratio of expression levels of a given protein between pairs of cell lines was plotted against the normalized ratio of the abundance of *in vitro* substrates on the given protein between the same pair of cell lines. The Pearson correlation coefficients are shown on the plots.

### PWM for *in vitro* substrates obtained from different cell lines

Next, we evaluated the predictability of kinase substrates based on PWM. For this evaluation, only phosphorylated serine substrates were included, because the number of phosphorylated threonine substrates was small. *In vitro* substrates among different cell lines were classified into three classes (abundance ratio ≥2, 2>abundance ratio ≥1/2, abundance ratio <1/2) based on protein abundance ratio (Fig. S2a), and for each class, 400 or 1000 *in vitro* substrates were randomly selected as training sets to generate PWMs (Fig. S2b, Table S3). ROC curve analysis was performed to evaluate the kinase substrate predictability of PWMs derived from each class. As a test set, we randomly selected 200 known substrates for each kinase from the public database PhosphoSitePlus (positive data set), and combined them with 200 serine peripheral sequences randomly selected from the human proteome (negative data set). For these test sets, PWM scores were calculated using the PWM of each class. The results showed that PWMs generated from each class had similar AUCs (Fig. 2a, S3a, S3c). This trend was confirmed for all kinases and cell line pairs. These results indicate that the prediction of kinase substrates based on PWMs is not affected by differences in *in vitro* substrate abundance between cell lines. This trend was also confirmed using a training set consisting of different numbers of *in vitro* substrates (Fig. 2a, S3a, S3c). Furthermore, increasing the number of *in vitro* substrates in the training set improved the predictability of kinase substrates (Fig. 2b, S3b).

**Fig. 2.**
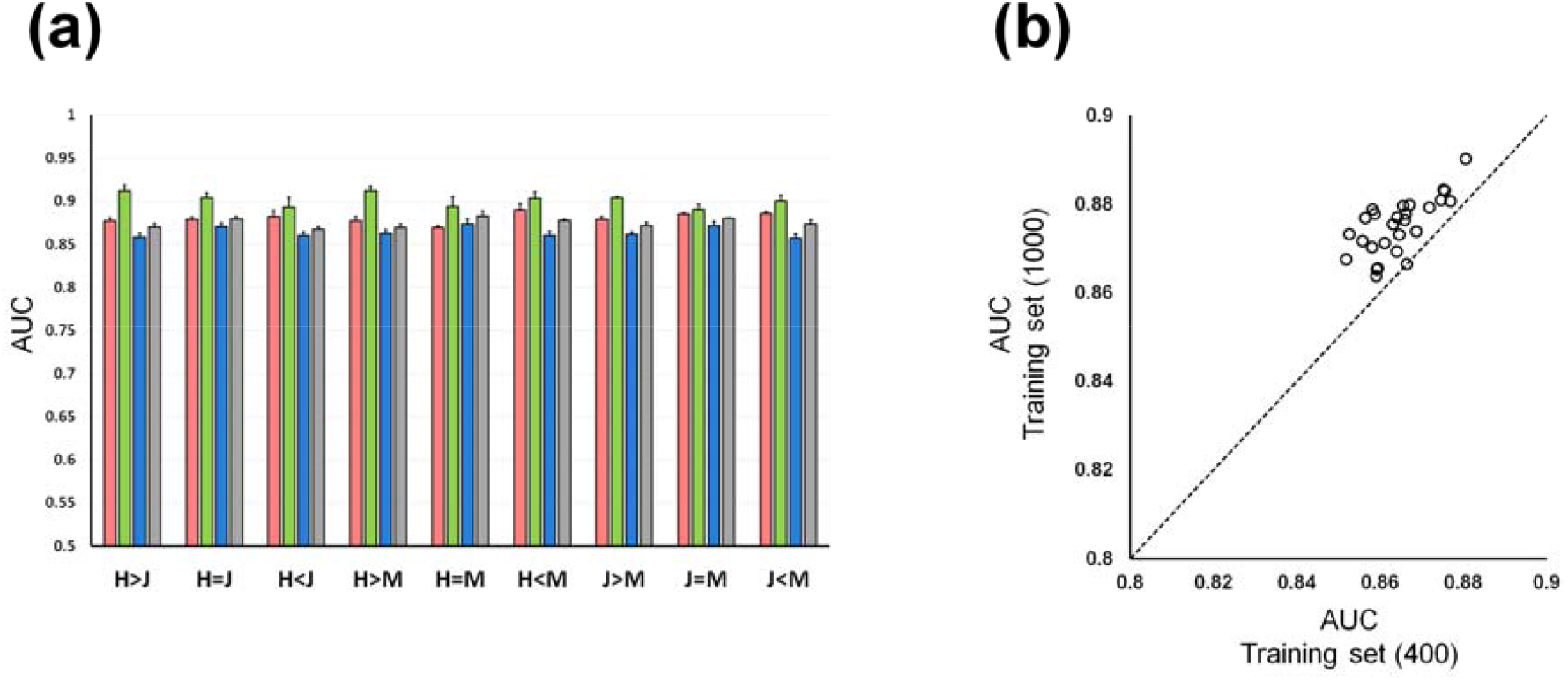
ROC curve analysis for prediction of kinase substrates with each cell-line-derived PWM. **(a) Kinase substrate prediction performance with each cell-derived PWM** The performance of kinase substrate prediction for each class was compared in Test 1. HeLa is indicated as H, Jurkat as J, and MCF7 as M. (Red) CK2 (Green) ERK 1 (Blue) PKA (from 400 *in vitro* substrates) (Gray) PKA (from 1000 *in vitro* substrates) **(b) Effect of *In vitro* PKA substrate number in the training set on kinase substrate predictions in Test 1** 27 training set pairs were evaluated using the same test set.

Finally, we evaluated the predictability of kinase substrates based on PWMs obtained from random sequence peptide arrays (PL-PWM) (Johnson et al., 2023), compared with those obtained from the cell extracts. As a result, PL-PWMs and cell extract-based PWMs provided similar predictability of kinase substrates for the three kinases (Fig. 3a). Furthermore, the correlations of PMW scores between replicates using the same cell line and between different cell lines, as well as the correlation of PMW scores between a random sequence peptide library and cell-extract substrate pools, were similar (Fig. 3b). These results indicate that the phosphorylation motifs obtained from the human cell-extracted substrate pool do not contain any specific bias arising from the starting material.

**Fig. 3.**
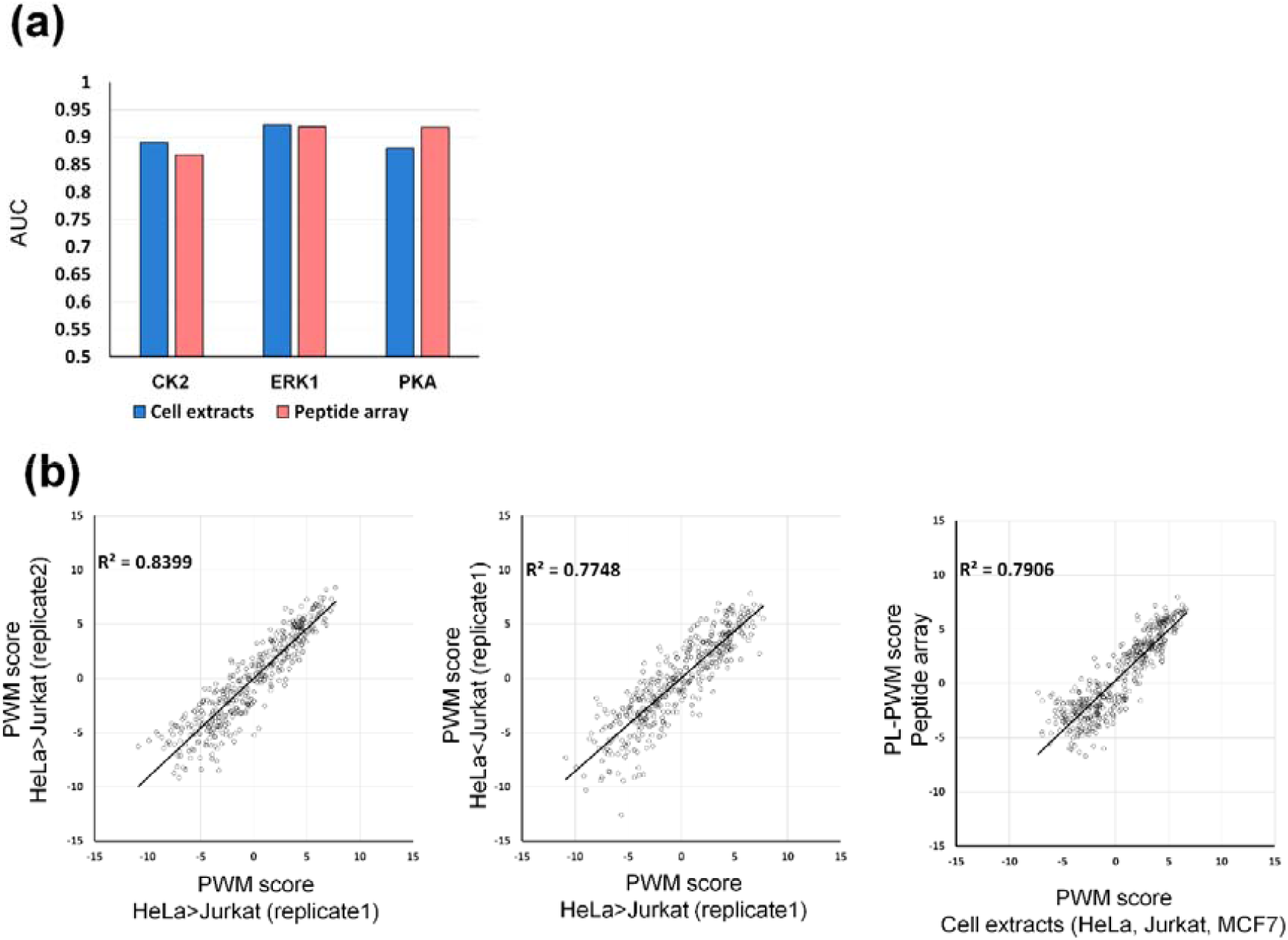
Evaluation of the predictability of kinase substrates using PWM score and PL-PWM score. **(a) ROC curve analysis for prediction of kinase substrates with PWM score and PL-PWM score** Comparison of the performance of kinase substrate prediction for each class in Test 1. **(b) Correlation between PWM score and PL-PWM score for ERK1 in Test 1**

## Discussion

The first large substrate pools employed for phosphorylation motif extraction were synthetic peptide libraries, as used for example in NetPhorest (Miller et al., 2008). When using peptide libraries, peptides of different sequences should ideally be immobilized individually in each well, but in practice, due to cost and labor considerations, a mixture of peptides of different sequences, identical only in one amino acid in one location, is immobilized in each well. As an alternative, cell lysate can be used as a substrate pool, and after phosphorylation reaction, the peptides are identified and quantified by LC/MS/MS using phosphoproteomics techniques, which efficiently provide information on each individual peptide. In this study, we demonstrated that in the latter method, the use of cell-extracted substrate pools obtained from cell lines with different proteome profiles does not affect the phosphorylation motifs extracted. We also compared the results of this method with those in a recently published study using random sequence peptide arrays (Johnson et al., 2023) and showed that both methods have comparable kinase-substrate predictability. On the other hand, unlike peptide libraries, cell-extracted substrate pools do not include artificial sequences. However, it would be possible to synthesize the desired peptides and mix them with the cell-extracted substrate pool. In any case, cell-extracted substrate pools are easier to prepare and more cost-effective than synthetic peptide pools. Our results suggest that this method will be applicable to kinases with other phosphorylation motifs, and we anticipate that cell extracts can continue to be used as substrate pools for profiling phosphorylation motifs.

## Experimental procedures

### Materials

Fetal bovine serum and BCA protein assay kit were purchased from Thermo Fisher Scientific. Amicon Ultra 10K was purchased from Merck Millipore. TSAP and sequencing-grade modified trypsin were purchased from Promega. Recombinant kinases were purchased from Carna Biosciences. All other chemicals were obtained from FUJIFILM Wako, unless otherwise specified.

### Cell culture

HeLa cells from cervical carcinoma and MCF7 cells from mammary carcinoma were cultured in DMEM containing 10% fetal bovine serum and 100 μg/mL penicillin/streptomycin. Jurkat cells from acute T-cell leukemia were cultured in RPMI1640 containing 10% fetal bovine serum and 100 μg/mL penicillin/streptomycin. All cells were maintained in an incubator at 37 °C under humidified 5% CO_2_ in air.

### *In vitro* kinase assay using cell extracts

*In vitro* kinase reaction was performed with recombinant kinases and cell extracts as described previously (Niinae et al., 2021). In brief, cells were washed and harvested with ice-cold PBS. Proteins were extracted from cells with phase-transfer surfactant (Masuda et al., 2008), and the buffer was replaced with 40 mM Tris-HCl (pH7.5) by ultrafiltration using an Amicon Ultra 10K at 14,000 g. Protein amount was determined with a BCA protein assay kit and the solution was divided into aliquots containing 100 μg. For each cell line, the following procedures were repeated in triplicate experiments. Proteins were reacted with 1 μL of TSAP, and TSAP was inactivated by heating to 75 °C for 1 hr. For *in vitro* kinase reaction, proteins were reacted with 1 μL recombinant kinase or water at 37 °C in the reaction buffer (40 mM Tris-HCl pH 7.5, 20 mM MgCl_2_, 1 mM ATP) for 3 hr. Then, proteins were reduced with 10 mM DTT, alkylated with 50 mM IAA, diluted 2-fold with 50 mM ABC buffer and digested with Lys-C (w/w 1:100) and trypsin (w/w 1:100). Phosphopeptides were enriched from the tryptic peptides with aliphatic hydroxy acid-modified metal oxide chromatography (HAMMOC) (Sugiyama et al., 2007), desalted using SDB-XC StageTips, and suspended in the loading buffer (0.5% TFA and 4% ACN) for nanoLC/MS/MS analyses.

### Global proteome analyses of cell extracts

Aliquots containing proteins were reduced with 10 mM DTT, alkylated with 50 mM IAA, diluted 2-fold with 50 mM ABC buffer and digested with Lys-C (w/w 1:100) and trypsin (w/w 1:100). Then, peptides were desalted using SDB-XC StageTips, fractionated at basic pH (Rappsilber et al., 2007) and suspended in the loading buffer (0.5% TFA and 4% ACN) for LC/MS/MS analyses.

### NanoLC/MS/MS analyses

NanoLC/MS/MS analyses were performed on an Orbitrap Fusion Lumos (Thermo Fisher Scientific) connected to an Ultimate 3000 pump (Thermo Fisher Scientific) and an HTC-PAL autosampler (CTC Analytics, Zwingen, Switzerland). Peptides were separated on a self-pulled needle column (150 mm length x 100 μm ID, 6 μm opening) packed with Reprosil-C18 AQ 3 μm reversed-phase material (Dr. Maisch, Ammerbuch, Germany). The flow rate was set to 500 nL/min. The mobile phase consisted of (A) 0.5% acetic acid and (B) 0.5% acetic acid in 80% acetonitrile. Three-step linear gradients of 5-10% B in 5 min, 10-40% B in 100 min for global proteome analysis or in 60 min for phosphopeptides obtained by *in vitro* kinase reaction, and 40-99% B in 5 min were employed. The MS scan range was *m/z* 300-1500. MS scans were performed by the Orbitrap with *r* = 120,000 and subsequent MS/MS scans were performed by the Orbitrap with *r* = 15,000. Auto gain control for MS was set to 4.00 × 10^5^ and that for MS/MS was set to 5.00 × 10^4^. The HCD was set to 30.

### Database searching

For all experiments, the raw MS data files were analyzed by MaxQuant v1.6.17.0 (Cox & Mann, 2008). Peptides and proteins were identified by means of automated database searching using Andromeda against the human Swiss-Prot Database (version 2020-08, 20,368 protein entries) with a precursor mass tolerance of 20 ppm for the first search and 4.5 ppm for the main search and a fragment ion mass tolerance of 20 ppm. The enzyme was set as Trypsin/P with two missed cleavages allowed. Cysteine carbamidomethylation was set as a fixed modification. Methionine oxidation and acetylation on protein N-term were set as variable modifications. For phosphopeptides, phosphorylations on serine, threonine and tyrosine were also set as variable modifications. Match-between runs were performed. The search results were filtered with FDR < 1% at the peptide spectrum match (PSM) and protein levels. Phosphosites were filtered to accept only those for which the localization score was > 0.75.

### Data analysis

The peak area of each peptide in MS1 was quantified using MaxQuant (Cox & Mann, 2008). For the following analysis of phosphoproteome data, phosphorylation sites quantified in the control sample were excluded from the substrate list of the corresponding kinase-treated sample and imputed as intensity zero. Also, the leading protein out of Leading proteins was used and only quantitative values derived from monophosphorylated peptides were used. For the following analysis of global proteome data, the leading protein out of Majority protein IDs was used.

### PWM score

The probability of observing residue *x* in position *i*, from *in vitro* substrates for CK2, ERK1 or PKA is computed as follows:

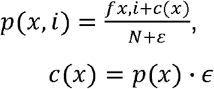

Where *fx, i* is the frequency of observing residue *x* at position *i* and *N* is the total number of sequences. *c(x)* is a pseudo count function which is computed as the probability of observing residue b in the human proteome based on the human Swiss-Prot Database (version 2017-04, 20,199 protein entries), *p(x)*, multiplied by, defined as the square root of the total number of sequences used to train the position weight matrix (PWM). This avoids infinite values when computing logarithms. Probabilities are then converted to weights as follows:

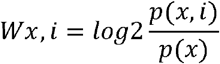

where *p(x) =* background probability of amino acid *x; p(x, i)* = corrected probability of amino acid *x* in position *i*; *Wx, i =* PWM value of amino acid *x* in position *i*. Given a sequence *q* of length *l*, a score *PWM score* is then computed by summing log_2_ weights:

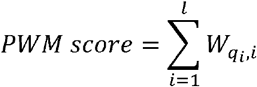

where q_*i*_ is the *i*^th^ residue of *q*. In this study, the score was computed using the flanking ±7 residues surrounding each phosphosite, except for comparison with random sequence peptide array data (Johnson et al., 2023). In the comparison with random sequence peptide array data, the score was computed using the flanking -5∼+4 residues surrounding each phosphosite.

For the PWM score based on the random sequence peptide library (PL-PWM score), PWM (called PSSM in the original paper (Johnson et al., 2023)) scores of the three kinases were extracted from the supplementary table (“ser_thr_all_norm_scaled_matrice”) in the report of Johnson et al. (2023).

PL-PWM score was calculated as follows:

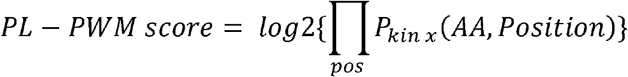

### ROC analysis

To evaluate the kinase substrate prediction performance of PWMs, ROC analysis was performed using 200 known substrates of a given kinase extracted randomly from PhosphoSitePlus (Hornbeck et al., 2015) as positive test set and 200 sequences around a serine residue extracted randomly from the human Swiss-Prot Database as negative test set. Two test sets (Test 1, Test 2) were created and used for duplicate analyses. All ROC analyses were carried out using the ROCR package in R (Sing et al., 2005).

PWM was generated using 400 or 1000 *in vitro* substrates extracted randomly from *in vitro* substrates grouped into three classes, or from total *in vitro* substrates derived from the three cell lines. In all evaluations, there were no overlaps of sequence between the positive test set, negative test set and training data set.

## Data Availability

The mass spectrometry proteomics data have been deposited at a public repository of ProteomeXchange Consortium (http://proteomecentral.proteomexchange.org) via the jPOST partner repository (https://jpostdb.org)(Okuda et al., 2017) with the data set identifier PXD042871.

## Author contributions

T.N., N.S. and Y.I. designed the research. T.N. performed the research and analyzed the data. T.N. and Y.I. wrote the paper.

## Acknowledgements

We would like to thank members of the Department of Molecular Systems BioAnalysis for fruitful discussions. This work was supported by JST Strategic Basic Research Program CREST (No. JPMJCR1862), AMED Advanced Research and Development Programs for Medical Innovation CREST (JP18gm1010010), and JSPS Grants-in-Aid for Scientific Research No. 17H03605, 21H02459, 23H04924 to Y. I., 18H04799, 20H04845, 21H02466 to N. S. and 21J15068 to T. N.

## Notes

### Competing Interest Statement

The authors have declared no competing interest.

### Summary of Updates

Figure 2, 3 revised

## References

Bekker-Jensen, D.B., Bernhardt, O.M., Hogrebe, A., Martinez-Val, A., Verbeke, L., Gandhi, T., Kelstrup, C.D., Reiter, L., & Olsen, J.V. (2020). Rapid and site-specific deep phosphoproteome profiling by data-independent acquisition without the need for spectral libraries. Nat. Commun. 11, 787.

Bodenmiller, B., Wanka, S., Kraft, C., et al. (2010). Phosphoproteomic analysis reveals interconnected system-wide responses to perturbations of kinases and phosphatases in yeast. Sci. Signal. 3, rs4.

Cox, J., & Mann, M. (2008). MaxQuant enables high peptide identification rates, individualized p.p.b.-range mass accuracies and proteome-wide protein quantification. Nat. Biotechnol. 26, 1367–1372.

Dinkel, H., Chica, C., Via, A., Gould, C.M., Jensen, L.J., Gibson, T.J., & Diella, F. (2011). Phospho.ELM: a database of phosphorylation sites--update 2011. Nucleic Acids Res. 39, D261–D267.

Geiger, T., Wehner, A., Schaab, C., Cox, J., & Mann, M. (2012). Comparative proteomic analysis of eleven common cell lines reveals ubiquitous but varying expression of most proteins. Mol. Cell. Proteomics 11, M111.014050.

Hornbeck, P.V., Zhang, B., Murray, B., Kornhauser, J.M., Latham, V., & Skrzypek, E. (2015). PhosphoSitePlus, 2014: mutations, PTMs and recalibrations. Nucleic Acids Res. 43, D512–D520.

Huang, S.-Y., Tsai, M.-L., Chen, G.-Y., Wu, C.-J., & Chen, S.-H. (2007). A systematic MS-based approach for identifying in vitro substrates of PKA and PKG in rat uteri. J. Proteome Res. 6, 2674–2684.

Humphrey, S.J., Azimifar, S.B., & Mann, M. (2015). High-throughput phosphoproteomics reveals in vivo insulin signaling dynamics. Nat. Biotechnol. 33, 990–995.

Hutti, J.E., Jarrell, E.T., Chang, J.D., Abbott, D.W., Storz, P., Toker, A., Cantley, L.C., & Turk, B.E. (2004). A rapid method for determining protein kinase phosphorylation specificity. Nat. Methods 1, 27–29.

Imami, K., Sugiyama, N., Imamura, H., Wakabayashi, M., Tomita, M., Taniguchi, M., Ueno, T., Toi, M., & Ishihama, Y. (2012). Temporal profiling of lapatinib-suppressed phosphorylation signals in EGFR/HER2 pathways. Mol. Cell. Proteomics 11, 1741–1757.

Imamura, H., Sugiyama, N., Wakabayashi, M., & Ishihama, Y. (2014). Large-scale identification of phosphorylation sites for profiling protein kinase selectivity. J. Proteome Res. 13, 3410–3419.

Imamura, H., Wagih, O., Niinae, T., Sugiyama, N., Beltrao, P., & Ishihama, Y. (2017). Identifications of Putative PKA Substrates with Quantitative Phosphoproteomics and Primary-Sequence-Based Scoring. J. Proteome Res. 16, 1825–1830.

Inamori, K., Kyo, M., Nishiya, Y., Inoue, Y., Sonoda, T., Kinoshita, E., Koike, T., & Katayama, Y. (2005). Detection and quantification of on-chip phosphorylated peptides by surface plasmon resonance imaging techniques using a phosphate capture molecule. Anal. Chem. 77, 3979–3985.

Inamori, K., Kyo, M., Matsukawa, K., Inoue, Y., Sonoda, T., Tatematsu, K., Tanizawa, K., Mori, T., & Katayama, Y. (2008). Optimal surface chemistry for peptide immobilization in on-chip phosphorylation analysis. Anal. Chem. 80, 643–650.

Johnson, J.L., Yaron, T.M., Huntsman, E.M., et al. (2023). An atlas of substrate specificities for the human serine/threonine kinome. Nature 613, 759–766.

Keshava Prasad, T.S., Goel, R., Kandasamy, K., et al. (2009). Human Protein Reference Database--2009 update. Nucleic Acids Res. 37, D767–D772.

Masuda, T., Tomita, M., & Ishihama, Y. (2008). Phase transfer surfactant-aided trypsin digestion for membrane proteome analysis. J. Proteome Res. 7, 731–740.

Miller, M.L., Jensen, L.J., Diella, F., et al. (2008). Linear motif atlas for phosphorylation-dependent signaling. Sci. Signal. 1, ra2.

Needham, E.J., Parker, B.L., Burykin, T., James, D.E., & Humphrey, S.J. (2019). Illuminating the dark phosphoproteome. Sci. Signal. 12.

Niinae, T., Imami, K., Sugiyama, N., & Ishihama, Y. (2021). Identification of Endogenous Kinase Substrates by Proximity Labeling Combined with Kinase Perturbation and Phosphorylation Motifs. Mol. Cell. Proteomics 20, 100119.

Okuda, S., Watanabe, Y., Moriya, Y., Kawano, S., Yamamoto, T., Matsumoto, M., Takami, T., Kobayashi, D., Araki, N., Yoshizawa, A.C., Tabata, T., Sugiyama, N., Goto, S., & Ishihama, Y. (2017). jPOSTrepo: an international standard data repository for proteomes. Nucleic Acids Res. 45, D1107–D1111.

Pan, C., Olsen, J.V., Daub, H., & Mann, M. (2009). Global effects of kinase inhibitors on signaling networks revealed by quantitative phosphoproteomics. Mol. Cell. Proteomics 8, 2796–2808.

Rappsilber, J., Mann, M., & Ishihama, Y. (2007). Protocol for micro-purification, enrichment, pre-fractionation and storage of peptides for proteomics using StageTips. Nat. Protoc. 2, 1896–1906.

Sing, T., Sander, O., Beerenwinkel, N., & Lengauer, T. (2005). ROCR: visualizing classifier performance in R. Bioinformatics 21, 3940–3941.

Sugiyama, N., Masuda, T., Shinoda, K., Nakamura, A., Tomita, M., & Ishihama, Y. (2007). Phosphopeptide enrichment by aliphatic hydroxy acid-modified metal oxide chromatography for nano-LC-MS/MS in proteomics applications. Mol. Cell. Proteomics 6, 1103–1109.

Sugiyama, N., Imamura, H., & Ishihama, Y. (2019). Large-scale Discovery of Substrates of the Human Kinome. Sci. Rep. 9, 10503.

Wasserman, W.W., & Sandelin, A. (2004). Applied bioinformatics for the identification of regulatory elements. Nat. Rev. Genet. 5, 276–287.

